# Top-down and bottom-up interactions rely on nested brain oscillations

**DOI:** 10.1101/2024.05.23.595462

**Authors:** Jelena Trajkovic, Domenica Veniero, Simon Hanslmayr, Satu Palva, Gabriela Cruz, Vincenzo Romei, Gregor Thut

**Affiliations:** Department of Cognitive Neuroscience, Faculty of Psychology and Neuroscience, Maastricht University, 6229 ER, Netherlands; School of Psychology, University of Nottingham, Nottingham, NG7 2RD, UK; Centre for Cognitive Neuroimaging, School of Psychology and Neuroscience, University of Glasgow, Glasgow, G12 8QB, UK; Centro studi e ricerche in Neuroscienze Cognitive, Dipartimento di Psicologia, Alma Mater Studiorum – Università di Bologna, Campus di Cesena, Cesena, 47521, Italy; Facultad de Lenguas y Educación, Universidad Antonio de Nebrija, Madrid, 28015, Spain

## Abstract

Adaptive visual processing is enabled through the dynamic interplay between top-down and bottom-up (feedback/feedforward) information exchange, presumably propagated through brain oscillations. Here we causally tested for the oscillatory mechanisms governing this interaction in the visual system. Using concurrent TMS-EEG, we emulated top-down signals by a single TMS-pulse over the Frontal Eye Field (right-FEF), while manipulating the strength of sensory input through the presentation of moving concentric gratings (compared to a control-TMS site). FEF-TMS without sensory input led to a top-down controlled occipital phase-realignment, alongside higher fronto-occipital phase-connectivity, in the alpha/beta-band. Sensory input in the absence of FEF-TMS increased occipital gamma activity. Crucially, testing the interaction between top-down and bottom-up processes (FEF-TMS during sensory input) revealed an increased nesting of the bottom-up gamma activity in the alpha/beta-band cycles. This establishes a causal link between phase-to-power coupling and top-down modulation of feedforward signals, providing novel mechanistic insights into how attention interacts with sensory input at the neural level, shaping rhythmic sampling.

## Introduction

Flexible and adaptive perception is made possible by top-down (e.g. predictive) regulation of sensory input. This involves a large-scale brain network, in which top-down signals influence lower-level visual areas (1–4). For example, when attending to a specific spatial location, top-down attentional control enables enhanced processing of information in the attentional spotlight (5); with the Frontal Eye Field (FEF) being a key area for this control (6). Specifically, non-human primate work has demonstrated that task-related attentional signals generated in the FEF exert top-down influence on visual areas (7–9), and that micro-stimulation of FEF yields attention-like effects in the visual system (10,11). Similarly, a causal role of the FEF in attentional control has been demonstrated in human participants, in whom the activation of FEF by means of non-invasive transcranial magnetic stimulation (TMS) causes changes in visual cortex activity (12–14) and perception (15–17).

At the same time, brain oscillations have been identified to play a role in attention and perception, whereby attention-controlled oscillatory synchronization in specific frequency bands both within and between brain regions is thought to facilitate visual processing (18–20). For instance, non-human primate studies found that tasks emphasizing top-down processing evoke stronger lower frequency (alpha/beta) synchronization, whilst feed-forward processing is associated with high-frequency (gamma) oscillations (7,21,22). In support of this dichotomy, micro-stimulation of pathways and cortical layers involved in feedback or feedforward signalling is leading to stronger synchronization at alpha/beta or gamma frequencies respectively (22–26). Similarly, in the human visual system inter-areal rhythmic influences are observed to predominate in the alpha/beta-bands along feedback projections (18,27), but in the gamma-band along feedforward projections (19). This accords with earlier findings revealing that sensory processing in primary visual cortices is related to gamma activity (28–31), with this activity being modulated by both low-level physical features (such as stimulus size and spatial location (32)) and attentional demands (33,34). Finally, causal probes of top-down signalling through FEF-stimulation by TMS in human participants – mirroring previous micro-stimulation probes in animals (10,23) – point to top-down control being enabled by frequency-specific oscillatory changes between distant brain areas at alpha/beta-frequency, namely their phase re-alignment (35).

Taken together, the above findings suggest that top-down control from higher-order areas is implemented through oscillatory processes at alpha/beta activity, while sensory input is processed and forwarded through fast oscillatory activity at gamma frequency. However, how these top-down rhythms interact with bottom-up brain signals is still unknown. One candidate mechanism is cross-frequency interactions, where the slower (alpha/beta-frequency) oscillations organize gamma-activity through phase-amplitude coupling such that gamma-bursts become nested within the alpha/beta phase, thereby enabling a coordinated interplay between neural elements of a large-scale attention network (36,37). By extension, phase-amplitude coupling (PAC) of alpha/beta to gamma oscillations could enable a coupling between feedback– and feedforward streams for optimal performance. Yet, causal tests of such a mechanism of top-down control over bottom-up input are lacking.

The aim of the present study was to causally probe this interaction in the human brain using concurrent TMS-EEG. Specifically, we stimulated FEF through a TMS pulse causing top-down influences in the alpha/beta-range over lower-level visual areas (replicating (35)), while manipulating sensory input in the form of a continuously presented moving sine-grating. By simultaneously recording electroencephalography (EEG), we tested whether the FEF-stimulation reorganised the sensory-driven gamma-activity to become phase-coupled to the top-down generated alpha/beta-cycles as an expression of top-down interactions over bottom-up signals.

## Results

In healthy human volunteers (N=30), we employed brief TMS pulses to activate an attention control centre in the prefrontal cortex (FEF-TMS), compared to a control stimulation site (M1foot-TMS). At the same time, we manipulated the strength of sensory input through the presentation of moving concentric gratings (GRATING +/− condition) that lasted for 5 seconds. The participants were instructed to respond with a buttonpress everytime they perceived a pause (glitch) in the motion of the grating (see Fig 1). The duration of the glitch was thresholded for each participant (for more details, see Methods). We then tracked the neural expressions of the associated top-down and bottom-up signals with concurrent EEG.

**Fig 1.**
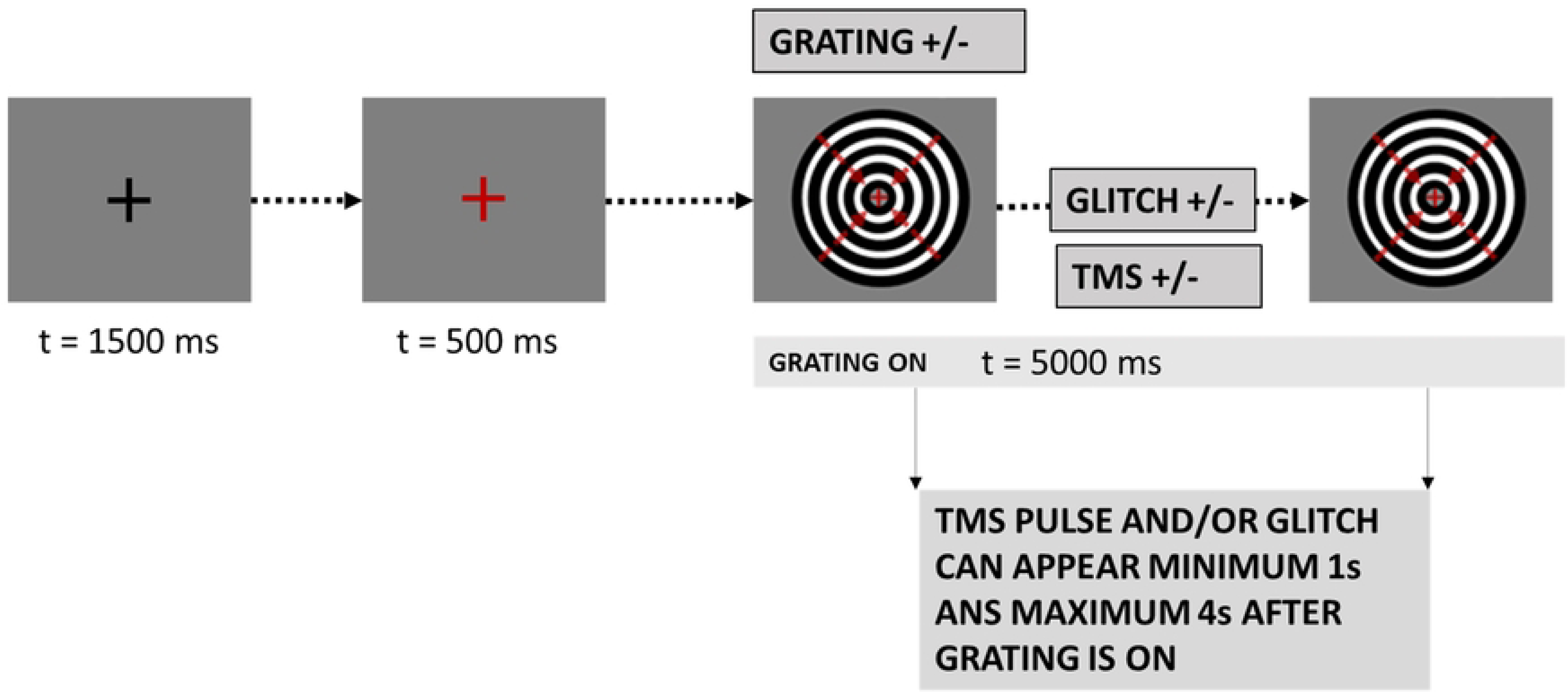
*Experimental and task design*. Trial Sequence. Each trial began with a black fixation cross lasting 1500 ms. Afterwards, the cross turned red indicating the beginning of the trial. After 500ms, the visual grating stimulus was presented. It consisted of a black and white sine-grating contracting inwards and was displayed for 5 seconds. In the grating absent condition, a red fixation cross was displayed at the same time. Some grating-present trials were followed by a target glitch in the grating motion, and participants had to respond with a button press whenever they perceived the glitch. Moreover, there were 3 TMS conditions: right FEF-TMS, control TMS over the right foot area, and no TMS. Both the TMS pulse and the target glitch could appear from 1 second earliest and to 4 seconds latest after grating onset.

### Sensory input induces gamma-band activity

The first analysis aimed at confirming that our experimental manipulation of sensory input had the expected consequences, i.e., that the moving grating enhanced gamma activity over posterior sites. To do so, all trials with versus without the moving grating (GRATING+/−) were compared in a period before any TMS-pulse and the occurrence of possible motion glitches (first second after moving grating onset). The permutation-based analysis confirmed that there is significantly higher gamma synchronization in trials with the grating stimulus present as compared to the no-grating condition, lasting for the whole post-grating time window analyzed (0-800 ms) (Fig 2A, but also see Supporting Figure 1 for the same analysis on the whole grating duration: 0-5000 ms). These differences span both low and higher gamma frequency bands (50-80 Hz). Importantly, differences in map topography indicate that these effects were constrained to posterior electrodes (see Fig 2B). As expected, grating presentation was also accompanied by a desynchronization across lower 10-20 Hz (alpha-beta) frequencies.

**Fig 2.**
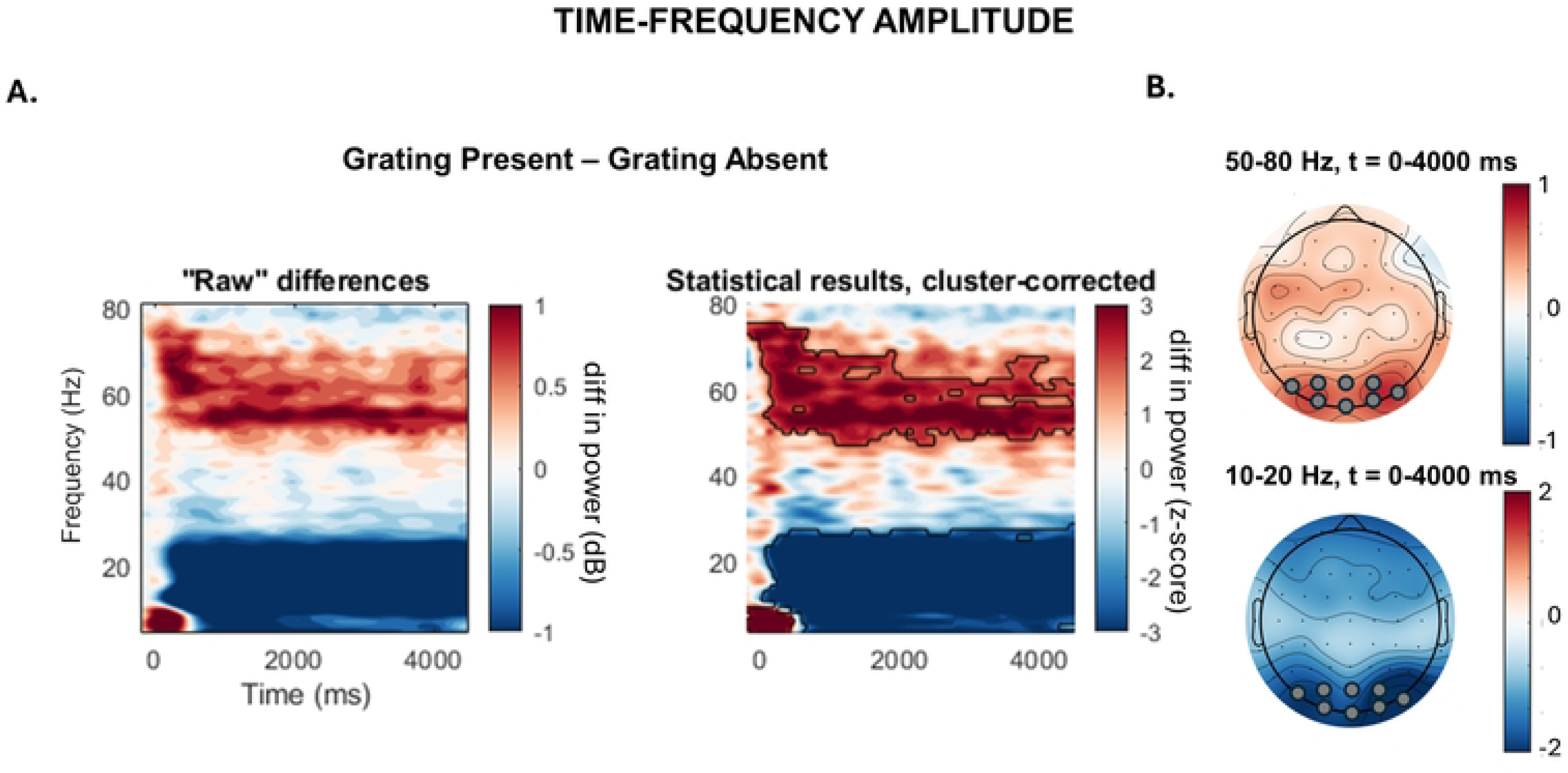
*Time-Frequency Analysis:* Grating effects. (**A**) Left panel: Power differences in time-frequency plots of the posterior electrodes (O2, O1, POz, Oz, PO8, PO7, PO4, PO3) between the grating present and grating absent condition (GLITCH+/−). The frequency range for the analysis (y-axis) is from 5-80Hz. The time range for the analysis (x-axis) is from –200 to 800 ms, where 0 is the time point of the grating onset. Right panel: Z-scores of the permutation-based analysis between grating present and grating absent condition. Significant clusters are framed with the black line. (**B**) Topographies of the significant clusters of the amplitude differences in the gamma (upper map) and higher alpha/lower beta frequency bands (lower map). Electrodes used for the time-frequency analysis are marked with grey circles. diff = difference; dB = decibel; Hz=hertz; t=time.

### FEF exerts top-down control over posterior areas through phase-realignment and enhanced connectivity in alpha/beta frequency bands

We investigated the influence of FEF-TMS on posterior activity via analysis of intertrial phase coherence (ITPC), in comparison to a control TMS site (TMS of nearby M1-foot area). Based on a previous report (35), we expected FEF-activation by TMS to lead to a posterior phase-reset of oscillatory activity in the alpha/beta frequency band. Here, we replicate these findings. In trials without a grating stimulus (GRATING-), there was a significantly higher ITPC for FEF-TMS vs control-TMS in the lower beta frequency band (13-20 Hz) over posterior sites in the right (stimulated) hemisphere, until around 400ms after the TMS pulse (Fig 3A). In analogy, for trials with a grating stimulus (GRATING+), FEF-TMS, relative to control-TMS, led to higher ITPC over right posterior sites in a similar lower beta frequency range, including also higher alpha-band activity (11-19Hz), for mainly the first 250ms after the TMS pulse (Fig 3B). There were no significant effects across the left posterior electrodes, in either grating present or absent condition.

**Fig 3.**
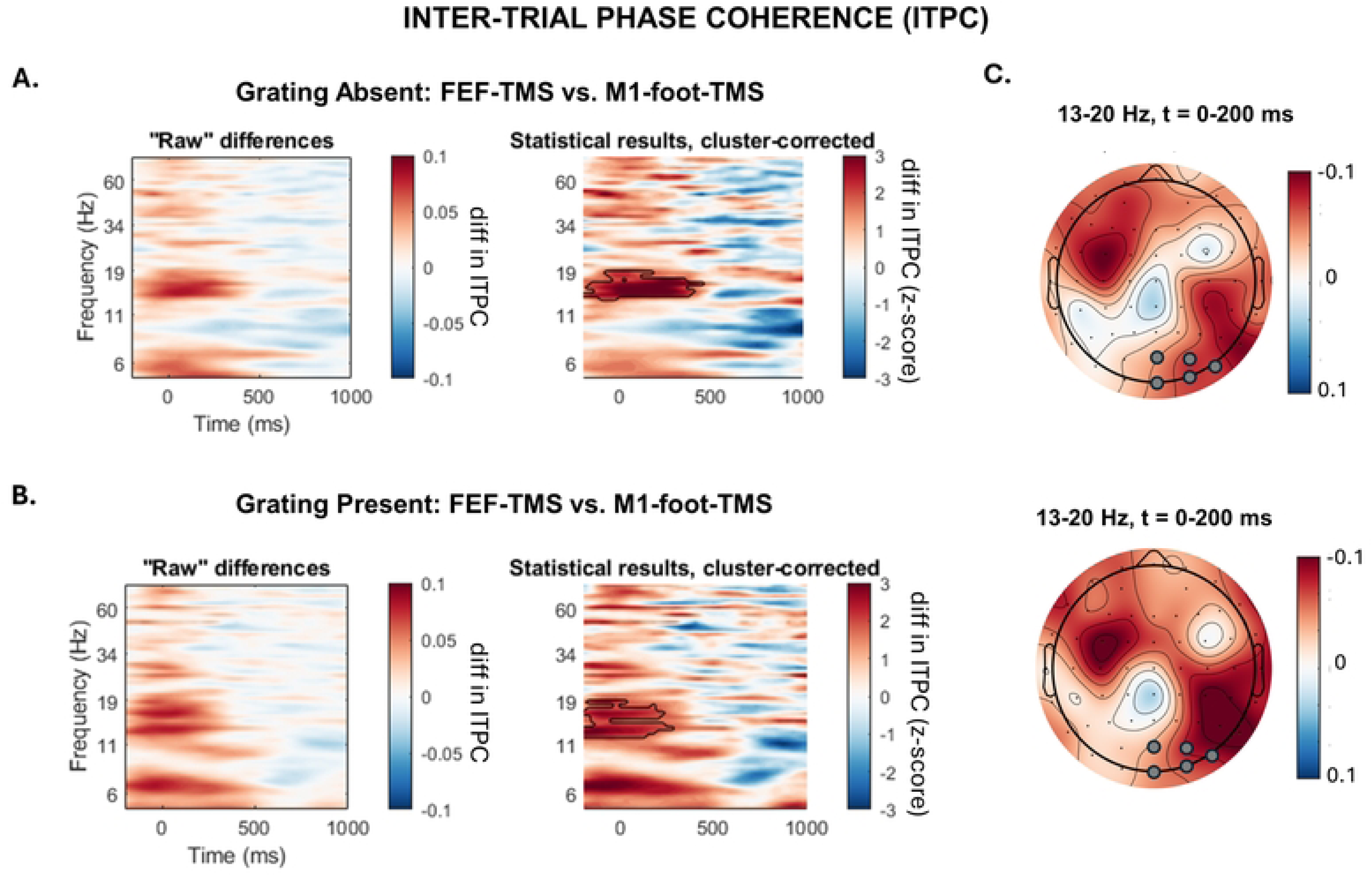
ITPC Analysis: Differences between FEF-TMS and control TMS (over M1foot). (**A**) **Grating Absent condition.** Left panel: Inter-trial phase coherence (ITPC) differences of the posterior cluster in the stimulated (right) hemisphere (electrodes: O2, POz, Oz, PO8, PO4) between FEF-TMS and control TMS. The time range for the analysis (x-axis) is from –200 to 1000 ms, where 0 point is the timing of the TMS-pulse. The frequency (y-axis) is from 5-80 Hz. Right panel: Z-scores of the permutation-based statistical analysis between FEF-TMS and control stimulation (M1foot-TMS). Significant clusters are framed with black lines. (**B**) **Grating Present condition.** Left panel: Raw differences in ITPC plots of the posterior cluster in the stimulated (right) hemisphere (electrodes: O2, POz, Oz, PO8, PO4) between FEF-TMS and control stimulation (M1foot-TMS). The frequency range for the analysis (y-axis) is from 5-80 Hz. The time range for the analysis (x-axis) is from –200 to 1000 ms, where 0 point is the timing of the TMS-pulse. Right panel: Z-scores of the permutation-based analysis between FEF-TMS and control stimulation (M1foot-TMS). **C.** Topographies of the significant clusters of the identified ITPC differences in the grating absent (upper map) and grating present condition (lower map). Electrodes used for the ITPC analysis are marked with grey circles. diff = difference; ITPC = inter-trial phase coherence; Hz=hertz; t=time.

To assess if FEF-TMS (vs foot-TMS) enhances inter-areal phase coupling between frontal and posterior sites, we calculated the weighted phase lag index (wPLI): a phase lag-based measure not affected by volume conduction. Subsequently, non-parametric permutation analysis was used to compare wPLI values across the two stimulation sites (FEF-TMS, M1-foot) in the two grating stimulus conditions (GRATING+ and GRATING-). Connectivity was estimated in the 300ms window following TMS, between frontal and parieto-occipital electrodes, both in the TMS-stimulated and non-stimulated hemisphere, in the frequency-band identified as significant via ITPC. The results indicate that there is enhanced fronto-posterior connectivity for FEF-TMS as compared to control-TMS. Importantly, these differences were only significant for fronto-posterior electrode pairs in the right (stimulated) hemisphere (Fig 4A). As expected, these differences were visible both in the grating stimulus present (Fig 4C) and absent (Fig 4B) condition, as they likely represent a top-down control mechanism that should be present independently of sensory input.

**Fig 4.**
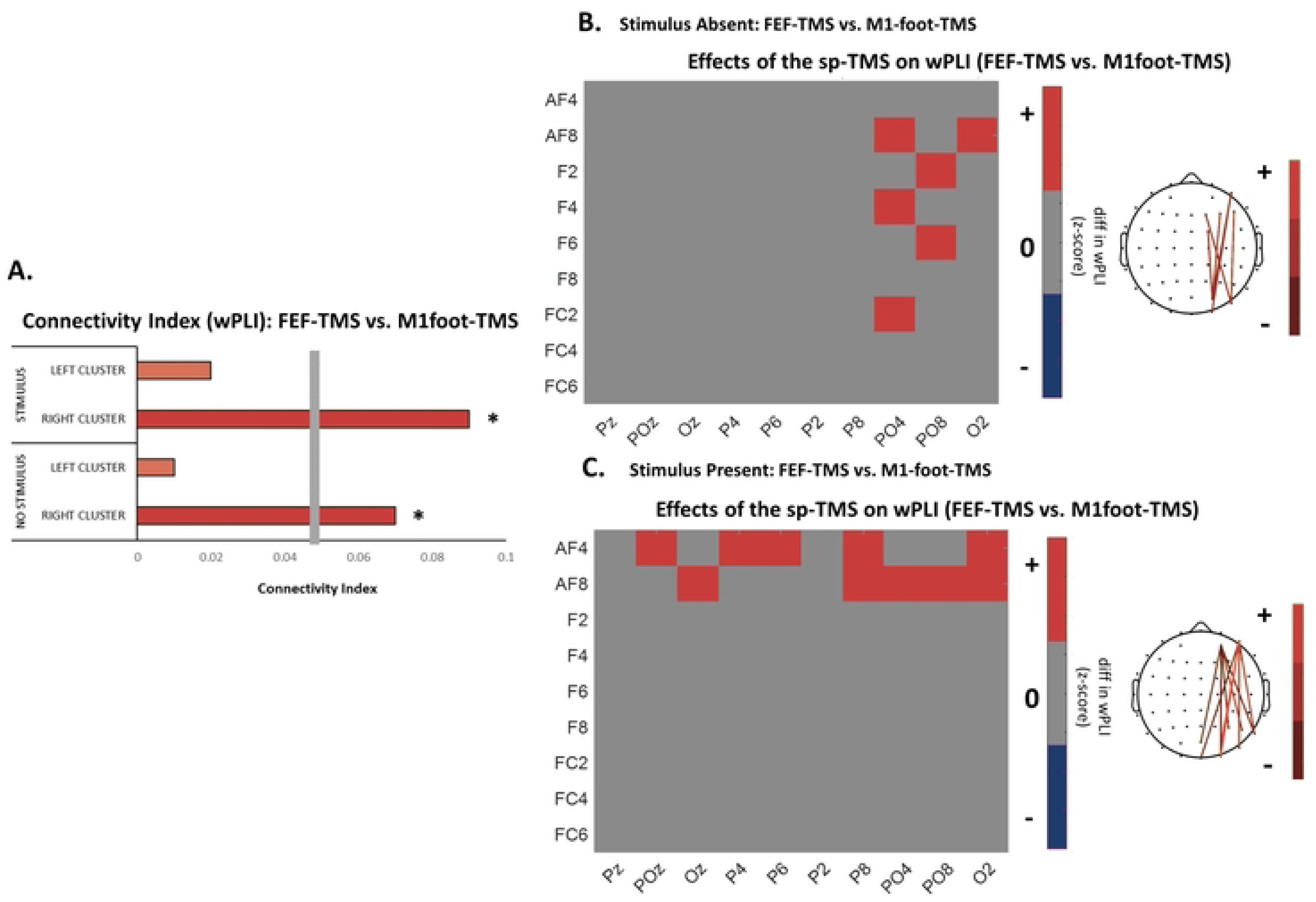
Changes in interregional coupling between fronto-posterior electrode pairs when contrasting activity recorded during FEF-TMS and control-TMS (over M1foot). (**A**) Representation of the connectivity index for each of the experimental conditions (grating present vs absent) and the two hemispheres (right, stimulated and the left, non-stimulated hemisphere); grey vertical bar shows the statistical threshold. Red ink indicates increases in interregional coupling for the FEF-TMS vs, control stimulation. (**B**) **Grating absent Condition.** Left: Connectivity matrix measured by the weighted phase lag index (wPLI) across all electrodes included in the regions of interest for the grating absent condition in the higher alpha/lower beta range (13-20 Hz), in the right (stimulated) hemisphere. Red ink indicates significant increases in interregional coupling for the FEF-TMS vs. control stimulation. Right: Topographical representations of the electrodes showing significantly increased coupling for the FEF-TMS vs, control TMS (**C**) **Grating Present Condition.** Left: Connectivity matrix measured by the weighted phase lag index (wPLI) across all electrodes included in the regions of interest for the grating present condition in higher alpha/lower beta range (13-20 Hz), in the right (stimulated) hemisphere. Red ink indicates significant increases in interregional coupling for the FEF-TMS vs. control stimulation. Right: Topographical representations of the electrodes showing significantly increased coupling for the FEF-TMS vs. control TMS. wPLI = weighted phase-lag index.

### Top-down control and sensory processing interact through cross-frequency coupling

So far, we have demonstrated gamma-band activity over posterior sites to be related to bottom-up signalling of sensory input and/or sensory processing, as well as fronto-to-posterior phase realignment and connectivity enhancement – both in the alpha/beta band – to reflect top-down influences. We next tested whether the top-down signal may structure bottom-up input through phase-amplitude coupling, potentially enabling the integration of these two mechanisms of information processing. If so, we would expect gamma activity related to sensory input to be nested in distinct phases of the lower (alpha/beta) frequency, that is the bottom-up input to be top-down phase-controlled by FEF-generated signals.

To investigate this hypothesis, we calculated the modulation index of phase-amplitude coupling for the 200ms following the TMS pulse, between low-frequency and high-frequency activity (frequency ranges tested: 5-20Hz vs. 30-80Hz), across right parieto-occipital sites. Our results revealed an increase of phase-amplitude coupling over posterior electrodes when FEF is activated by TMS (vs control TMS over the M1-foot area) for the case when the stimulus was present (GRATING+, Fig 5B). More specifically, gamma amplitude (from 60-90Hz) was coupled to the lower beta phase (at 13-20 Hz), with a higher gamma activity for the bins around the beta peak (Fig 5C and 5D). Crucially, these differences between the two TMS protocols were not present in the absence of sensory input (GRATING-, Fig 5A), which shows that both top-down and bottom-up signals have to be present for phase-amplitude coupling to occur. This suggests that alpha/beta to gamma phase-amplitude coupling is indeed a mechanism that allows both processes to interface.

**Fig 5.**
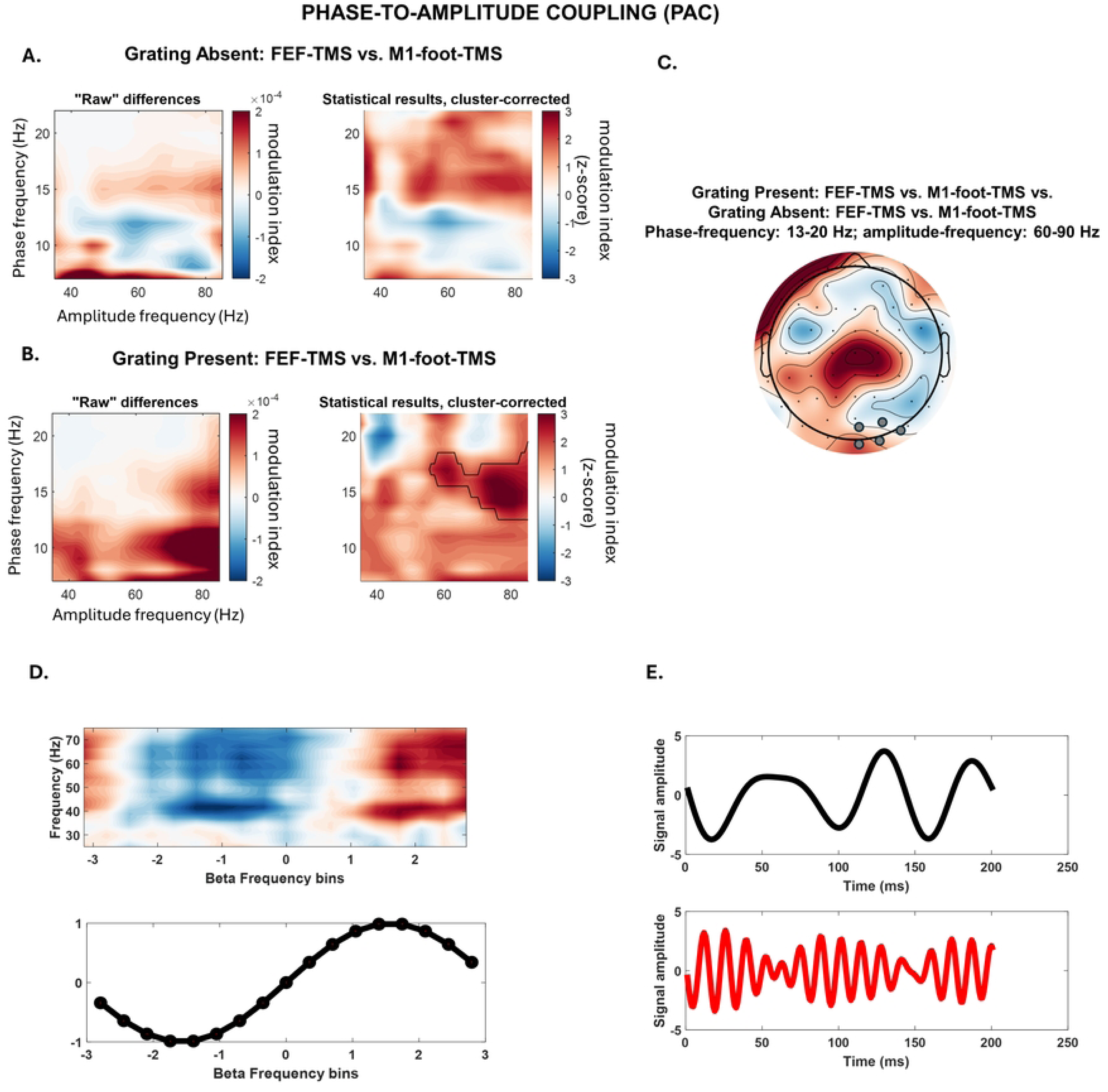
Phase-to-amplitude Coupling (Modulation Index) after the TMS pulse (0-200 ms). (**A**) **Grating Absent condition.** Left panel: Raw differences in modulation index (MI) plots of the posterior cluster in the stimulated (right) hemisphere (electrodes: O2, POz, Oz, PO8, PO4) between FEF-TMS and control stimulation (M1foot-TMS). The frequency for lower phase frequency is shown in the y-axis. The frequency for the higher gamma amplitude-in the x-axis. Right panel: Z-scores of the permutation-based analysis between FEF-TMS and control stimulation (M1foot-TMS). No significant clusters were identified. (**B**) **Grating Present condition.** Left panel: Raw differences in modulation index (MI) plots of the posterior cluster in the stimulated (right) hemisphere (electrodes: O2, POz, Oz, PO8, PO4) between FEF-TMS and control stimulation (M1foot-TMS). The frequency range for the analysis of the lower phase-frequency (y-axis) is from 5-20 Hz. The frequency range for the analysis for the higher gamma amplitude-frequency (x-axis) is from 30-80 Hz. Right panel: Z-scores of the permutation-based analysis between FEF-TMS and control stimulation (M1foot-TMS). Significant clusters are framed with the black line. (**C**) Topographies of the significant clusters of the identified MI differences in the grating present vs. grating absent condition. Electrodes used for the MI analysis are marked with grey circles. (**D**) Beta (15Hz) phase sorted gamma power is shown for all the data for the experimental condition that yielded significant modulation index (FEF-TMS when stimulus is present). Note the higher gamma activity for the bins around the beta peak. (**E**) Data for the single trial from the experimental condition that yielded significant modulation index (FEF-TMS when grating is present) filtered in the gamma and beta frequency. Note that gamma increases mostly in beta peaks. Hz=hertz; t=time.

## Discussion

Perception is much more than the passive reception of sensory input: it is the result of the complex interactions of internal signals based on priors with the incoming sensory input (38,39). Brain oscillations may enable this interplay, where feedback and feedforward processing is carried by slower and faster oscillatory activity, respectively, and where these two streams of processing may interact through cross-frequency mechanisms (22–25,40,41). Yet, a causal test of how feedback routes the sensory-driven signal through oscillatory mechanisms is lacking. Here we tested whether the generation of a top-down signal causes visually-driven, high-frequency activity in the posterior cortex to be nested to the phase of top-down generated low-frequency oscillations. To this end, we emulated an attentional impulse via FEF-stimulation through a TMS pulse in the presence versus absence of sensory activity, while examining effects on EEG.

Our results first confirmed that a continuous visual input, in the form of a moving sine-grating (42), results in synchronization of gamma-band activity across the posterior brain areas, corroborating prior evidence that the feedforward drive of sensory input is accompanied by gamma synchronization (28–31). This gamma increase is thought to enhance synaptic summation (19), thereby boosting the transfer of information at-hand (43). Second, when emulating an attentional impulse by FEF-TMS (vs TMS over a control area), we observed a high alpha/low beta oscillatory phase-realignment and enhanced connectivity between the frontal and posterior EEG sensors both when sensory input was present and absent. This is in line with numerous studies indicating that alpha/beta activity reflects top-down processes (7–9) and confirms the notion that reorganization of alpha/beta-phase, through long-range connectivity in the same frequency band, may represent a basic mechanism for feedback communication (35). Third, in the same posterior cluster, and when FEF-TMS was combined with sensory input, we observed gamma activity to be locked to the phase of the alpha/beta frequency (as compared to TMS over the control site), but not when no sensory input was present. Importantly, the coupled low/high frequency oscillations exactly matched in frequency the alpha/beta– and gamma-range identified as significantly associated with top-down and bottom-up sensory processing in isolation. This pattern of results therefore establishes a causal link between cross-frequency coupling and top-down modulation of feedforward signals, and corroborates that this top-down versus bottom-up integration is achieved through phase-to-amplitude cross-frequency coupling.

The alpha-oscillation (and adjacent brain rhythms) has not only been identified as governing feedback processing streams but has also been shown to be perceptually relevant. More specifically, its phase across visual areas has been related to perceptual outcome, where threshold stimuli are sometimes perceived and sometimes missed based on the instantaneous phase at the moment of stimulus presentation (44–49). Accordingly, it has been proposed that these fluctuations reflect the cyclic changes of brain excitability indicating the rapid waxing and waning of visual and/or attention sampling, predicting perceptual outcome (45,47). While research on visual/attentional sampling has mostly pointed to effects in the alpha– or theta-range (50–53), it is conceivable that the phase-reorganization in the high alpha/low beta-band found here reflects the expression of another rhythmic sampling mechanism at the interface between attention and sensory processing, that comes about through feedback-controlled periodic modulation of feedforward signaling. In support of this view, we recently found that activation of the FEF via a single TMS pulse not only phase-resets posterior alpha/beta-oscillations but induces cyclic modulations of visual perception and extrastriate visual cortex (V5) excitability at high alpha/low beta-frequency, as tested by motion discrimination accuracy and moving phosphene perception (35). Interestingly, it has recently been proposed that attentional sampling is organized via two alternating attentional states (54,55). While one attentional state is thought to be related to alpha-band activity and the rhythmic (pulsed) suppression of visual processing, the second state, associated with better stimulus detection, is characterized by increased posterior gamma-activity, alongside enhanced beta-activity in FEF (55). According to this model, gamma-activity enables enhanced sensory processing at the cued location, while beta-activity leads to a decreased likelihood of shifting away from the attentional locus (54,55). In line with this model, our data suggest that beta-gamma interplay is indeed a fingerprint of FEF-related visual attention processes, that seems to shape rhythmic sampling at high alpha/low beta-frequency through frequency-specific feedback-over-feedforward interactions. It is an open question whether the low beta-rhythmicity observed here and in our previous study (35) is linked to motion processing, given that the employed experimental tasks share involvement of V5. It is conceivable that other task designs involving static visual stimuli or probing other nodes of the dorsal attentional network (e.g., parietal brain areas) will yield different oscillatory signatures, possibly in the alpha or theta range.

The framework positing that the communication between brain regions is based on nested oscillations (36,41) accommodates two prominent and related theories of oscillatory communication. The first one is the communication-through-coherence theory, where interregional communication is considered established when these regions oscillate at the same frequency with a stable phase-lag (56). The second one is the gating through inhibition hypothesis, where slower oscillations are considered to be associated with pulses of inhibition and as such can support interareal communication, through phase synchronization and release of inhibition (57,58). Finally, the unified framework based on nested oscillations posits that the communication between two regions is established by phase synchronization of oscillations at lower frequencies (<25 Hz), which serve as a temporal reference frame for information carried by high-frequency activity (>40 Hz) (36,41). Our results speak in favor of this theory, where top-down predictions from FEF are communicated through oscillatory coherence to the posterior cortex, where they dictate sensory information flow through the nesting of gamma oscillations.

In summary, by emulating attention signaling through FEF-TMS and manipulating bottom-up processing in human participants, the current study reveals phase-to-amplitude coupling as a causal mechanism of interactions between low-frequency top-down and high-frequency feedforward signals. Overall, our results corroborate the beta-gamma interplay as an important fingerprint of a FEF-related visual/attention sampling mechanism, in line with recent models and in addition to other, well-documented period sampling mechanisms at alpha and theta-frequency.

## Materials and Methods

### Participants

A total of thirty participants took part in the study (18 females; mean age ± SE = 24.6 ± 4.11 years). All participants had normal or corrected-to-normal vision and reported no contraindication to TMS (59) or any neurological, psychiatric, or relevant medical condition. All protocols were performed in accordance with ethical TMS standards and approved by the Ethical Committee of the College of Science and Engineering (University of Glasgow, 300210149). All participants gave written informed consent to participate in the study.

### Stimuli, task and procedure

#### Main Experimental Task

Participants were comfortably seated at a viewing distance of 57 cm from an LCD monitor (144 Hz refresh rate), in a dimly illuminated room, with their chin on a chinrest to ensure a stable head position. The main experimental session consisted of two TMS conditions during which EEG was continuously recorded: Single pulse TMS was applied to either the right Frontal Eye Field or a control area (right M1-foot area), while participants were presented with a moving visual grating (half of the trials, GRATING+) or no grating (other half of the trials, GRATING-). Data was collected in four separate blocks (2 blocks per stimulation site, pseudorandomized so that two consecutive blocks would not have the same stimulation site). In order to uphold the alertness of the participant, a visual task was implemented (see below).

Visual stimuli and tasks were created in Matlab (Psychtoolbox-3). In both TMS conditions (right FEF, right foot M1), the task structure, number of pulses and pulse timing were kept constant. Specifically, each trial began with a black fixation cross, which, after 1500ms, turned red to signal that the grating would shortly appear (see Figure 1 for trial structure). After 500 ms, either a continuous circular sine grating contracting inwards was presented for 5 seconds in the central visual field (GRATING+ condition), or no stimulus appeared (GRATING-condition). The task of the participant was to attend to the moving grating when present and to press a button whenever they noticed a glitch in the movement of the grating, while always keeping their eyes on the fixation cross. Motion glitches were implemented to ensure participants remained in an alert state and occurred in 1/3 to 1/2 of all GRATING+ trials (GLITCH+ condition), whilst in the rest of the GRATING+ trials, moving gratings were uninterrupted (GLITCH-condition). The duration of the glitch was individualized for each participant, during a separate titration session (see below), to ensure that the perception of the glitch was around the threshold. A TMS pulse was delivered on one of the two TMS sites (depending on the experimental block) in TMS+ conditions, following a jittered interval ranging from 1 to 4 s after the grating onset (or at the same times into a trial when no grating was presented). In trials where both the TMS pulse and the glitch were present (TMS+/GLITCH+), the timing of the TMS pulse and the GLITCH coincided. Finally, there were also trials without a TMS pulse (TMS-), to assess gamma activity during the time-window of possible TMS delivery (1-4 sec).

In total, the experiment consisted of 640 trials and lasted about 90 minutes. The design led to a maximum of 80 trials per condition to be used in the EEG analyses. Conditions included in the EEG analyses comprised FEF-TMS, control-TMS or no TMS (TMS+/−), with/without a stimulus (GRATING+/−) but no glitches (all GLITCH+ trials excluded for EEG analysis) (see Fig 6 for the details on trial number per condition).

**Fig 6.**
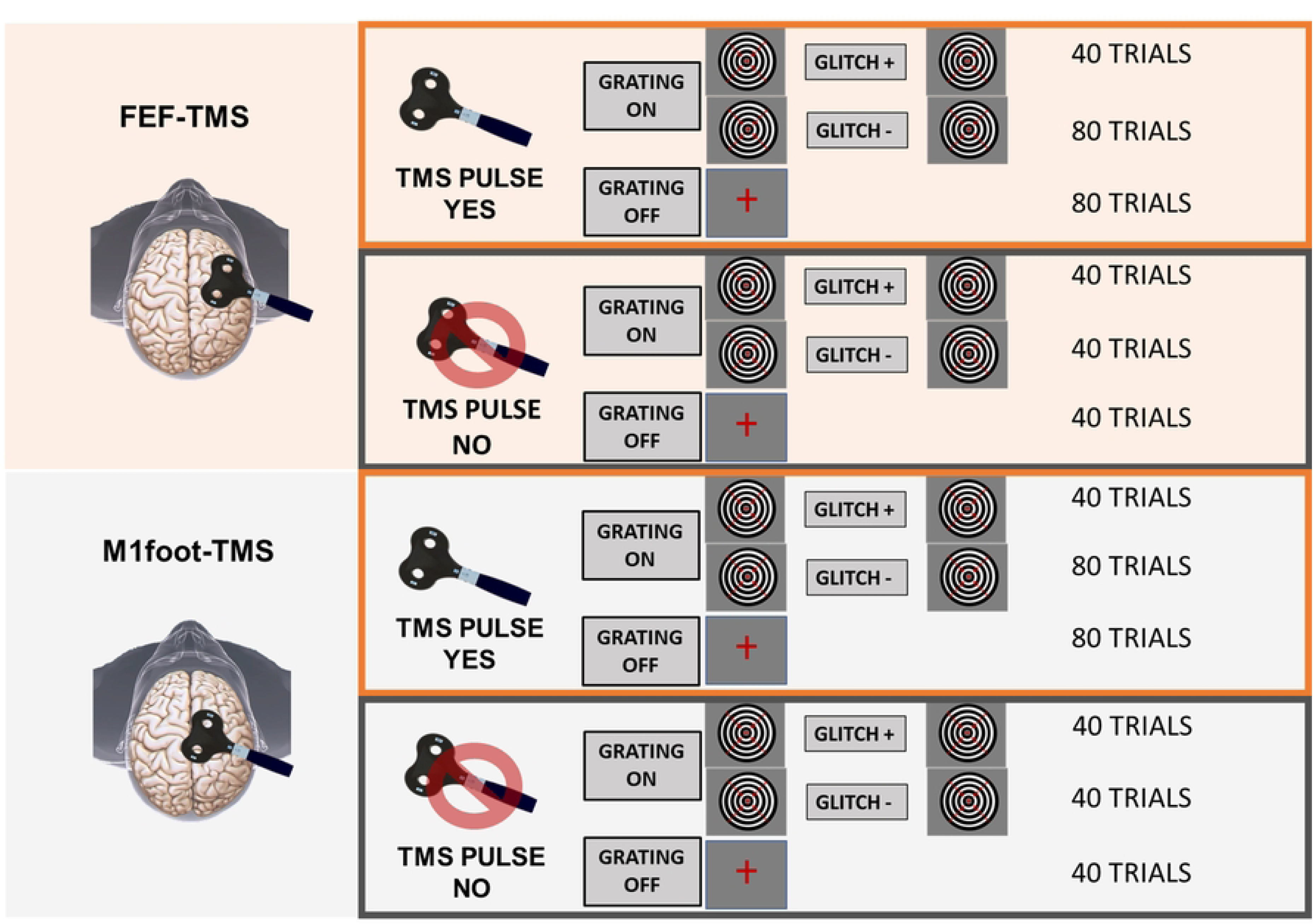
Experimental design. Task conditions with respective number of trials.

#### Titration Session

The duration of the glitch in the sine-grating movement, i.e., the number of frames during which the grating kept still, was individually thresholded for each participant via a blocked staircase procedure. Specifically, the titration session began with a glitch duration value of 20 frames (at 144Hz, 138.8ms) in an initial block (N = 10 trials), half of them being target and half of them catch trials (sine grating without the glitch). Subsequently, the percentage of correct trials was calculated and the glitch duration was adjusted accordingly (100% correct: glitch duration reduced by 6 frames; 95-86% correct: glitch duration reduced by 4 frames; 85-76% correct: glitch duration reduced by 2 frames; 75-56% correct: same glitch duration; 55-46% correct: glitch duration increased by 2 frames; 45-36% correct: glitch duration increased by 4 frames; 35-26% correct: glitch duration increased by 6 frames; 25-6% correct: glitch duration increased by 8 frames; 5–0% correct: glitch duration increased by 10 frames). The staircase procedure was run for 16 blocks in total (Ntotal = 160 trials), and glitch duration values for each block were fitted to a sigmoid function, to identify duration values corresponding to 70% of accurate glitch detection for each participant (mean glitch duration (SD) = 7.14 (2.12) frames), subsequently used in the experiment.

#### TMS

TMS was applied by means of a high-power Magstim Rapid2 machine (Magstim Company, Whitland, UK), with the stimulator connected to a 70mm standard figure-of-eight coil, triggered remotely using Matlab. TMS intensity was kept constant for the two TMS sites at 65% maximal stimulator output (MSO), based on prior evidence that TMS at this intensity effectively activates FEF (17,35,60). Likewise, the coil position was kept constant, placed tangentially with the handle of the coil pointing backwards and laterally approximately 45 degrees to the interhemispheric line (as in (35)).

To locate the target area, T1-weighted structural magnetic resonance images (MRI) were acquired via a 3T Siemens Trio Tim scanner (Siemens, Erlangen, Germany) per participant. Subsequently, the stimulation site for the right FEF was individually localized applying the Cortex-based Alignment (CBA) approach in Brain-Voyager QX 2.8 (Brain Innovation, Maastricht, The Netherlands), as in (35). Briefly, anatomical data was used to reconstruct the cortical surface of each participant, which was then aligned to an atlas brain that includes the probabilistic group maps of FEFs. These group maps were then back-transformed to the individual brain anatomy, with the TMS target defined as the centre of gravity for the area with the highest probability (61,62). FEF coordinates were on average (± SD) x=31.2 ± 3.7, y=−9.7 ± 2.9, z=49.74 ± 5.1. The control area (M1-foot area) was localized using averaged Talaraich coordinates identified in the literature (x=6.8, y=−14.8, z=69.8) (63). Finally, to target the right FEF and right M1-foot area, and to keep coil position and orientation constant, Talairach coordinates obtained with the CBA (and average coordinates for the control area) were imported to a frameless stereotactic neuro-navigation system (Brainsight; Rogue solution).

### TMS-EEG recordings – acquisition and pre-processing

EEG activity was recorded using a TMS-compatible EEG system (BrainAmp MRplus, BrainProducts), acquiring EEG from 61 TMS-compatible Ag/AgCl multitrode electrodes (EasyCap GmbH, Herrsching, Germany) mounted on a cap and positioned according to the 10– 10 International System. Additional electrodes were used as ground (TP9) and recording reference (AFz). The signal was acquired at a sampling rate of 5000 Hz, and bandpass filtered at 0.1–1000 Hz. Electrode impedance was maintained below 5 KΩ. An additional electrode was positioned on the outer canthus of the left eye to record eye movements.

TMS artefacts were identified and removed using an open-source EEGLab extension, the TMS-EEG signal analyzer (TESA)(64). First, EEG data were epoched around sine-grating stimulus appearance or TMS pulse onset (between –2000ms and 1000ms). In trials where TMS-pulse was not present, an additional marker in the TMS-timing window was added to allow for epoching. Afterwards, a baseline correction was applied by subtracting the average of the entire epoch from each data point (demeaning the data). Data around the TMS pulse (–1ms +10ms) was removed and replaced with zeros and cubic interpolation of the removed window was performed prior to down-sampling the data (from 5000Hz to 1000Hz). At this point, data were visually inspected to remove noisy trials. Interpolated data around the pulse was again removed prior to Individual Component Analysis (ICA). Specifically, a fast ICA algorithm was used (pop_tesa_fastica function) to identify individual components representing artefacts, along with automatic component classification (pop_tesa_compselect function), where each component was subsequently manually checked and reclassified when necessary. In this first round of ICA, only components with large amplitude artefacts, such as rhythmic-TMS-evoked scalp muscle artefacts, were eliminated (M_removedIC_±SD = 5.95 ± 1.83). Data around the TMS pulse were again interpolated (cubic interpolation) prior to applying pass-band (between 1 and 100Hz) and stop-band (between 48 and 52Hz) Butterworth filters. Subsequently, interpolated data were again removed prior to the second round of ICA, in order to remove all other artefacts, such as blinks, eye movement, persistent muscle activity and electrode noise (M_removedIC_±SD = 18.36 ± 5.95). Then, the TMS-pulse period was again interpolated (cubic interpolation), and data was re-referenced to the average of all electrodes. Finally, single trials were visually inspected and those containing residual TMS artefacts were removed. The described TMS artefact removal procedure was applied to all EEG data. On average, less than 15% of all epochs were removed (M = 14.19%, SD = 5.85%).

#### Time-Frequency analysis: Effects of VISUAL STIMULI (GRATING+/−) on oscillatory amplitude

As a first step, to confirm that the continuous sine grating did indeed lead to an increase of gamma activity, a time-frequency analysis was performed on the signal epoched around the grating stimulus onset. Specifically, complex Morlet wavelet convolution was applied to the signal, with morlet wavelet peak frequencies ranging from 3 to 80Hz in 60 logarithmic steps. The full width at half-maximum (FWHM) ranged from 500 to 200ms with increasing wavelet peak frequency. Subsequently, power values were normalized by decibel conversion, with the baseline period from –500ms to –200ms before grating stimulus onset. Finally, time-frequency differences in power between the trials with and without the sine-grating stimulus in parieto-occipital cluster (cluster electrodes: O2, PO4, PO8, Oz, POz, PO3, PO7, O1) were compared via cluster-based permutation testing. Specifically, z-scores for each individual data point in the TF plane were obtained by comparing the time-frequency map differences (trials with vs. without the sine-grating) with permuted (N=1000) random differences (obtained by randomizing the sign of difference map for every subject). Differences with z-values corresponding to p<0.05 were retained as significant. Subsequently, the results were cluster-corrected, whereas the cluster threshold was determined by the 5% biggest cluster size in the null distribution of permuted z-scores (thus p<.05).

#### Phase-realignment: Effects of FEF-TMS (vs control-TMS) on occipito-parietal activity

Next, we aimed to replicate the findings of Veniero et al. (2021) of enhanced beta phase-realignment in connected posterior areas after FEF activation by a TMS pulse. We used intertrial phase clustering (ITPC) (as in Veniero et al., 2021) to track the influence of FEF-stimulation on the posterior cortex across time-frequency space. ITPC measures the extent to which phase angles at each time-frequency-electrode point across the trials are nonuniformly distributed in polar space (Cohen, 2014). Phase angles were obtained by the same wavelet convolution procedure used for the time-frequency power analysis. Likewise, the same statistical permutation-based cluster analysis used for comparing amplitude differences was used for ITPC-difference plots between the two TMS protocols (FEF-TMS, M1-foot-TMS), separately for the stimulus present (GRATING+) and stimulus absent (GRATING-) conditions. The same posterior electrodes as for the time-frequency amplitude analysis were used, but separately for the right (O2, PO4, PO8, Oz, POz) and left electrode cluster (Oz, POz, PO3, PO7, O1), given the right-lateralized TMS over FEF/foot areas.

#### Connectivity changes: Effects of FEF-TMS (vs control-TMS) on fronto-posterior connectivity

Interareal connectivity was estimated in the sensor space between frontal and parieto-occipital electrodes of both hemispheres (extending the analyses of Veniero et al., 2021) via the weighted phase-lag index (wPLI) (66). This is a measure of phase lag-based connectivity, which accounts for non-zero phase lag/lead relations between two signal time series, as it defines connectivity as the absolute value of the average sign of phase angle differences and additionally deweighting vectors that are closer to the real axis such that those vectors have a smaller influence on the final connectivity estimate. By extension, this measure is insensitive to volume conduction and noise of a different origin, considered optimal for exploratory analysis as it minimizes type-I errors (Cohen, 2015). To obtain wPLI values, time series data was first transformed into the time-frequency domain via convolution with a family of complex Morlet wavelets (the number of cycles increased from 5 to 18 in logarithmic steps). Therefore, for frequencies ranging from 3 to 30Hz in 1Hz steps, first convolution by frequency-domain multiplication was performed and then the inverse Fourier transformation was taken. Phase was defined as the angle relative to the positive real axis, and phase differences were then computed between all possible pairs of electrodes. Finally, wPLI was calculated as the absolute value of the average sign of phase angle differences, whereby vectors closer to the real axis were de-weighted. Once wPLI values were extracted for every frequency bin (3 to 30Hz in 1Hz steps) and epoch, they were averaged to obtain distinct values for each experimental condition of interest (FEF and M1-foot TMS when sine-grating was present or absent) in the higher alpha/lower beta frequency band identified as significant via ITPC (13-20 Hz, see results). Subsequently, non-parametric permutation-based analysis (1000 iterations) was performed to compare the connectivity between the two TMS protocols, and to obtain phase connectivity difference maps of distinct electrode pairs. For these pairwise calculations of fronto-posterior connectivity, we included the following electrodes of the right (stimulated) hemisphere (frontal: AF4, AF8, F2, F4, F6, F8, FC2, FC4, FC6; posterior: Pz POz Oz P2 P4 P6 P8 PO4 PO8 O2), as well as homologous electrodes of the left hemisphere. Sensor differences with z-values corresponding to p<0.05 were retained as significant. The connectivity index of each condition was then estimated using the formula: CI = sig_pos/ sp_total, where sig_pos are connections that are significantly higher after FEF-TMS with respect to M1foot-TMS, and sp_tot are all possible connections (68,69). Furthermore, another permutation test was introduced to calculate the significance threshold for CI. Specifically, wPLI matrices of all the electrode pairs and experimental conditions were randomly permuted and compared 1000 times to obtain the distribution of randomly obtained differences in wPLI. Connectivity indices that exceeded a 95% confidence interval were considered statistically significant (connectivity threshold= 0.05).

#### Phase-Amplitude Coupling

Finally, the hypothesized interaction between the top-down signal emulated by the TMS pulse and feedforward sensory input was measured via phase-amplitude coupling, with the idea that the phase of low-frequency (alpha/beta) rhythms should modulate the amplitude of high-frequency (gamma) oscillations. Phase-amplitude coupling was quantified via the modulation index (70). Briefly, the signal was first filtered at the two frequency ranges under analysis (low frequencies = 5-20Hz with 4Hz bandwidth; high frequencies = 30-80Hz with 10Hz bandwidth). Then, the Hilbert transform was applied to obtain the phase time series for the lower frequency range, and the time series of the amplitude envelope for higher frequencies. Next, these phases were binned and the mean of amplitude over each bin was calculated and normalized by the sum over all the bins, resulting in mean amplitude distribution over phase bins. Finally, the modulation index (MI) was obtained by measuring the divergence of the observed amplitude distribution from the uniform distribution. MI was calculated for every cross-frequency point in time-window of interest (0-200 ms post-TMS), concatenated between trials, for the right parieto-occipital cluster (POZ, OZ, O2, PO4, PO8), identified as significant with amplitude and ITPC analyses. The same statistical permutation-based cluster analysis used for the amplitude and ITPC analyses was used for PAC-difference maps between two TMS sites, separately for grating-present and grating-absent conditions.

## Contributions

Conceptualization: GT, JT, SP, VR; Methodology: GT, SH, SP, GC, VR, JT, DV; Investigation: JT; Visualization: JT; Supervision: GT; Writing—original draft: GT, JT, GC, DV, VR, SH, SP; Writing—review & editing: GT, JT, GC, DV, VR, SH, SP

## Data Availability

All data reported in this paper have been made publicly available through the Open Science Framework: https://osf.io/9uhta/.

## Supporting Information

**S1 Figure.** *Time-Frequency Analysis for the whole grating duration*: Grating effects. **A.** Left panel: Raw differences in time-frequency plots of the posterior electrodes (O2, O1, POz, Oz, PO8, PO7, PO4, PO3) between grating present and grating absent condition (GRATING+/−). Frequency range for the analysis (y-axis) is from 5-80Hz. Time range for the analysis (x-axis) is from – 200 to 5000 ms, where 0 is the time point of the grating onset. Right panel: Z-scores of the permutation-based analysis between grating present and grating absent condition. Significant clusters are framed with the black line. **B.** Topographies of the significant clusters of the amplitude differences in the gamma frequency band (lower). diff = difference; dB = decibel; Hz=hertz; t=time.

